# Sphingomyelinase Licensing of Mesenchymal Stromal Cells Alters Lipid and Protein Metabolites for Immunomodulation

**DOI:** 10.1101/2025.07.17.665406

**Authors:** S’Dravious A. DeVeaux, Daniel C. Shah, Keji Rui, Nathan F. Chiappa, Hongmanlin Zhang, Nidhi Lal, Rhyland O’Neill, Young C. Jang, Luke Mortensen, Krishnendu Roy, Edward A. Botchwey

**Affiliations:** The Wallace H. Coulter Department of Biomedical Engineering, Georgia Tech and Emory, Atlanta, GA 30332, United States; Petit Institute of Bioengineering and Biosciences, Georgia Institute of Technology, Atlanta, GA 30332, United States; Regenerative Bioscience Center, Rhodes Center for ADS, University of Georgia, Athens, GA 30602, USA; School of Chemical, Materials and Biomedical Engineering, University of Georgia, Athens, GA, United States; School of Materials Science and Engineering, Georgia Institute of Technology, Atlanta, GA 30332, USA; Atlanta Veterans Affairs Medical Center, Decatur, GA, 30030, USA; School of Biological Science, Georgia Institute of Technology, Atlanta, GA 30332, USA; Department of Orthopaedics, Emory Musculoskeletal Institute, Emory University School of Medicine, Atlanta, GA 30329, USA; Department of Biomedical Engineering, School of Engineering, Vanderbilt University, Nashville, TN 37235, USA; Department of Pathology, Microbiology and Immunology, School of Medicine, Vanderbilt University, Nashville, TN 37235, USA; Department of Chemical and Biomolecular Engineering, School of Engineering, Vanderbilt University, Nashville, TN 37235, USA

## Abstract

Mesenchymal stromal cells (MSCs) are widely studied for their immunomodulatory and tissue reparative capabilities, but clinical translation has been hampered by inconsistent efficacy and limited standardization in manufacturing. While cytokine-based priming methods, such as interferon-gamma (IFN-γ) stimulation, have shown promise in enhancing MSC potency, alternative approaches targeting distinct biological metabolism integral to secretome and membrane architecture have not been explored in MSCs. In this study, we investigate sphingomyelinase (SMase), an enzyme that generates ceramide from sphingomyelin, as a novel lipid-based priming strategy to modulate MSC function. Here, human MSCs were treated with SMase and high-content imaging and morphological profiling revealed that SMase-treated cells adopted a phenotype overlapping with IFN-γ–licensed MSCs, including increased cell compactness and solidity. Lipidomic analysis showed broad alterations in sphingolipid species, and dynamic flux estimation (DFE) modeling predicted distinct metabolic shifts in SMase-treated cells compared to untreated controls. These changes were sustained up to 35 hours post-stimulation, indicating stable metabolic reprogramming. SMase priming also altered the MSC secretome, enriching for factors implicated in immune regulation. Functionally, SMase-primed MSCs retained the ability to suppress T-cell activation and promote anti-inflammatory macrophage phenotypes. Collectively, these findings demonstrate that SMase stimulation induces a durable, immunomodulatory-like state in MSCs through coordinated changes in lipid metabolism and secretory activity. This lipid-centric priming approach represents a promising alternative to cytokine-based licensing strategies and may support therapeutic MSC products.

## INTRODUCTION

Widely distributed across vascularized tissues, mesenchymal stromal cells (MSCs) are non-hematopoietic, multipotent progenitors with broad differentiation potential and significant therapeutic promise. They are defined by their plastic adherence, characteristic surface antigen expression, the ability to self-renew in vitro and differentiate into mesodermal lineages such as osteoblasts, chondrocytes, and adipocytes [1, 2]. MSCs have attracted significant interest due to their potent immunomodulatory and tissue reparative capabilities, largely mediated through the secretion of paracrine factors and extracellular vesicles [3–5]. These properties make MSCs a promising candidate for treating inflammatory and degenerative diseases [6]. Despite extensive preclinical success, the clinical translation of MSC-based therapies remains challenging. Major obstacles include inconsistencies in therapeutic outcomes, limited cell engraftment and persistence in vivo, and difficulties in standardizing manufacturing processes to meet quality standards [7, 8].

More recently, a landmark milestone in the field was achieved with the recent FDA approval of Remestemcel-L (Ryoncil), an allogeneic bone marrow-derived MSC therapy, for the treatment of steroid-refractory acute graft-versus-host disease (GVHD) in pediatric patients [9]. This approval represents a significant step forward in MSC-based therapeutics and highlights the importance of standardizing manufacturing protocols. As a means of standardization, the identification of critical quality attributes (CQAs) and critical process parameters (CPPs) has become a point of emphasis in MSC biomanufacturing. CQAs, such as the expression of immunomodulatory molecules (e.g., IDO, PGE2), cytokine secretion profiles, and extracellular vesicle composition are emerging as measurable indicators of functional efficacy [10–12]. Likewise, CPPs including culture media and preconditioning methods must be carefully controlled to ensure consistency and predictability across production batches.

Literature suggests that bioactive lipids, particularly sphingolipids, may play a role in modulating MSC behavior and potentially serve as novel CQAs and CPPs [13–16]. Sphingolipids are a class of bioactive lipids that have been shown to be involved in key cell signaling processes involved in proliferation, cell migration, membrane properties, and cell fate. To better understand the lipid landscape and their broad relationships, lipidomic profiling is now being explored as a method for evaluating MSC function and predicting in vivo therapeutic outcomes [17, 18]. Through these methods, sphingolipids have been found to be deeply involved in biological processes central to healing. For example, sphingosine-1-phosphate (S1P) and ceramide-1-phosphate (C1P) are elevated in damaged tissues and act as potent chemoattractants for MSCs and other regenerative cells [19, 20]. In addition to promoting homing and adhesion, sphingolipids are known to regulate angiogenesis, modulate macrophage polarization, and influence the secretion of pro-regenerative cytokines [21–24]. These roles position sphingolipids not only as biomarkers but as active agents in MSC-mediated wound healing and immune modulation. Among the enzymes that regulate sphingolipid metabolism, sphingomyelinase (SMase) has emerged as a promising modulator of MSC phenotype and function. SMase hydrolyzes sphingomyelin into ceramide, a key player implicated in stress response, apoptosis, and inflammation. Ceramide enrichment within the plasma membrane can influence MSC immune interactions and enhance the secretion of regulatory molecules [25]. Notably, SMase activity is elevated in inflamed tissues, suggesting its role in priming MSCs in situ [26–28].

Traditional exogenous licensing approaches in MSC research commonly involve stimulation with pro-inflammatory cytokines such as interferon-gamma (IFN-γ) and tumor necrosis factor-alpha (TNF-α) [29, 30]. These stimuli enhance the expression of key immunoregulatory molecules including indoleamine 2,3-dioxygenase (IDO), HLA-G, and PD-L1, thereby promoting MSC-mediated suppression of immune responses [31]. Lipid-based priming strategies, such as the use of SMase, present a complementary approach that may activate distinct membrane-associated signaling pathways and modulate cellular functions through mechanisms not directly targeted by cytokine exposure. For example, ceramide generation via SMase can reorganize membrane microdomains, alter intracellular trafficking, and impact metabolic state, potentially leading to broader or more durable changes in MSC phenotype [32–36]. Unlike cytokine priming, lipid modulation may also offer greater stability, scalability, and ease of integration into manufacturing workflows, making it a promising alternative or adjunct in MSC bioprocessing.

This study aims to explore the feasibility of using SMase as a novel priming agent to enhance the therapeutic potential of MSCs. Specifically, we investigate how SMase treatment alters cellular viability, modulates the lipidomic profile, secretome composition, and morphological features of MSCs, and whether these changes correlate with enhanced immunomodulatory function. We seek to determine whether lipid-driven modulation can serve as a viable and potential alternative for MSC licensed therapeutics. Furthermore, by linking biochemical signatures to functional outcomes, this work contributes to the growing effort to define more predictive and mechanistically grounded CQAs and CPPs for clinical-grade MSCs. In summary, leveraging lipid metabolism, and in particular SMase-driven pathways, represents a novel approach to improving MSC-based therapies. As the field shifts from empirical use to mechanism-based manufacturing, such strategies may pave the way for more consistent and effective cell therapies capable of meeting regulatory standards and clinical demands.

## MATERIALS AND METHODS

### MSC Culture Parameters and Treatment

BM-MSC donor RB183 was purchased from RoosterBio, Inc. (Frederick, MD) with a PDL of 8.9. These cells were considered as passage 0 (P0). The MSCs were thawed and expanded by protocols provided by RoosterBio, Inc. Cells were cultured in RoosterNourish™ media (High Performance Media Kit, KT-001) containing serum-derived RoosterBooster™ in T-225 flasks at an approximate density of 3,333 cells/cm^2^. Cells were grown to 80% confluence (3 – 4 days) and then harvested the following day per manufacturer protocol. Cells were lifted from the flasks by TrypLE Express (Invitrogen) and the enzyme was neutralized with spent media prior to collecting the cells by centrifugation. Cells were frozen in Cryostor CS5 cryopreservation media for future experiments and stored in the vapor phase of liquid nitrogen. Cells were frozen as P1. Prior to experimentation, cells were thawed and grown to ∼80% confluency before being treated with 0.3 U/mL of sphingomyelinase (S7651, Sigma-Aldrich, St. Louis, MO) for 24 hours. Cells for all experiments were P1-P3 and ∼12-18 PDL.

### Live/Dead Imaging

MSCs were cultured and treated as previously described. Viability was determined using LIVE/DEAD™ Viability/Cytotoxicity Kit (Invitrogen), according to the manufacture’s protocol. After incubating with the specified working reagents for 30 minutes at RT, the cells were visualized using a Zeiss 900 laser scanning confocal microscope (Zeiss) at 494nm and 528nm excitation.

### Alamar Blue Cell Viability

MSCs were seeded at 26,000 cells/cm^2^, allowed to be attached overnight and treated with SMase as previously described. After SMase treatment was removed, both groups were replenished with fresh RoosterNourish and 10% of alamarBlue® reagent (BUF012A, Bio-Rad Laboratories, Hercules, CA) and incubated for 4 hours. The absorbance was read at 570nm and 600nm. To calculate viability, the percent reduction formula was used provided by Bio-Rad protocol.

### Apoptosis Assay

MSCs were expanded into a T-225 flask for one passage (P2), before harvesting and seeding into a 6 well plate at a density of approximately 31,000 cells/cm^2^. MSCs were allowed to grow to 80-90% confluency before being treated with SMase. The SMase group was treated with 0.3 U/mL SMase for 24 hours. After 24 hours, MSCs were prepped for flow cytometry. MSCs were harvested and co-stained with Apotracker^TM^ Green (427403, BioLegend San Diego, CA) (400nM staining solution) a fluorophore that stains phosphatidylserine, and Zombie UV (423108, BioLegend) (1:1000). MSCs were analyzed using Cytek Aurora flow cytometer (Cytek Biosciences, Fremont, CA).

### Beta-Gal senescence Assay

MSCs were expanded into a T-225 flask for one passage (P2), before harvesting and seeding into a 12 well plate at a density of approximately 28,000 cells/cm^2^. Following the 24-hr SMase treatment, MSCs β-galactosidase were stained using the Senescence Detection Kit (CBA-230, Cell Biolab, San Diego, CA) protocol. In brief, MSCs were washed with PBS three times before fixing with fixation solution for 10 minutes at room temperature. The cells were washed three times with PBS, stained with working solution, and incubated overnight. To quantify the senescent cells, cells were counter stained with Hoechst 33342 (H3570, Thermo Fisher Scientific, Waltham, MA) following manufacturer’s protocol. MSC nuclei were imaged with the DAPI filter and senescent cells were imaged with brightfield microscopy, 3 images per well (n = 4 wells per group). To determine the percentage of senescent cells, ImageJ software maxima feature was used to obtain total cell count and senescent cell count [37]. The senescent cells were divided by the total cell count and averaged to find the percent senescent cell.

### Autophagosome Assay

MSCs were seeded onto µ-slides (80826, ibidi) at 10,000 cells/cm^2^ and allowed to attach overnight. Then were treated with SMase as previously described. Autophagy assay kit (ab139484, Abcam) was used for the assay. Media was removed and washed with 1X assay buffer. Dual detection reagent was added and allowed to incubate at 37°C for 30 minutes. 100 µl of 1X assay buffer was added back into the wells and the plate was analyzed using a BioTek Synergy H4 Hybrid plate reader at 480/530 nm for the green detection reagent and 340/480 nm for Hoechst. Prior to microscopy imaging, cells were fixed with 4% PFA and then washed thrice with the assay buffer. Images were acquired using a Nikon W1 Spinning Disk Confocal microscope equipped with a 40X objective. Five random fields of view (374.40 μm × 374.40 μm) were taken from each well for image analysis in Fiji [38]. Data from the five images per well were averaged to generate representative values for each well (n=4).

### Confocal Microscopy of Mitochondrial Networks

MSCs were seeded onto µ-slides (80826, ibidi) at 10,000 cells/cm^2^ and allowed to attach overnight. Then were treated with SMase as previously described. Following this treatment, MSCs were stained to assess mitochondrial morphology with MitoBrilliant™ Live 646 (7417, Tocris Bioscience) according to the manufacture’s recommendations. Briefly, stock staining solution was diluted to 200 nM in warm media, incubated with cells for 45 min at 37°C, and washed three times with warmed PBS for 5 min at 37°C. Cells were then fixed with 4% PFA for 10 min at 37°C, washed twice and incubated with Alexa Fluor™ Plus 555 Phalloidin (A34055, Invitrogen™) diluted 1:1000 for 60 min at RT. Hoechst 33342 (H1339, Invitrogen™) was diluted 1:2000 and incubated for 10 min at RT to visualize the nuclei.

Z-stack fluorescent images were acquired using a Nikon W1 Spinning Disk Confocal microscope using a 60X oil objective (1.4 numerical aperture, working distance of 0.13 mm). Five random fields of view (253.44 μm × 253.44 μm) were taken from each well for image analysis in Fiji [38]. Mitochondrial morphology and network characteristics were analyzed using the Mitochondria Analyzer plugin [39]. Data from the five images per well were averaged to generate representative values for each well (n=4).

### Lipid Metabolism Tracer

MSCs were seeded into a 24 well plate and cultured as described above. Once ∼80% confluent, cells were treated with SMase for 24 hours. The treatment was removed, and cells were washed with PBS^-/-^ then quickly incubated with 10 µM of C6 NBD-sphingomyelin (810218, Avanti Polar Lipids) for 2 hours. Cells were imaged using the GFP filter on a Lionheart FX automated microscope (Agilent BioTek, Santa Clara, CA). The cells and conditioned media were then collected for lipid extraction, as previously described [40]. Samples were resuspended in mobile phase (850:150:15 MeOH:H_2_O:H_2_PO_4_), analyzed using a Prominence High Performance Liquid Chromatography (HPLC) (Shimadzu UFLC, flow rate at 1mL min^-1^, excitation = 460 nm, emission = 535 nm) equipped with a C18 250 x 4.6 mm LC column (Phenomenex, Torrance, CA), and normalized to protein concentration. Total protein was quantified using the Pierce™ BCA Protein Assay Kit (23225, Thermo Scientific™) according to the manufacturer’s protocol.

### Lipid Metabolism Dynamic Flux Estimation Modeling

All computation and figure generation for DFE was done in MATLAB 2023b (Mathworks). First, cubic smoothing spline functions were fit to the dynamic concentration data using the csaps function. The value of the smoothing parameter was manually adjusted for each spline to capture the trend of the data without overfitting and was always between 0.6 and 0.9. Next, the derivatives of the cubic smoothing splines were computed using the fnder function. These derivatives at each time point, i, were used to construct a derivative vector as follows:

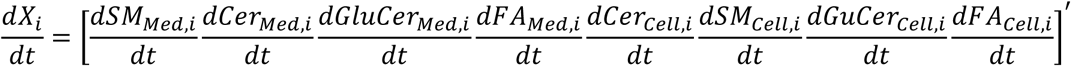

This vector was substituted into the left-hand side of the following flux balance equation.

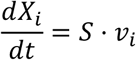

Here, S is the stoichiometric matrix and vi is the vector of fluxes at time point i. The stoichiometric matrix was set up based on the network structure shown previously. Given that the matrix is full row rank, there is only one unique solution to the flux balance equation which is calculated by inverting the matrix. This procedure is repeated for all time points to generate the full-time course of fluxes. The fluxes for the control and stimulated groups were then compared.

### Morphological Feature Imaging and Quantification

MSCs were seeded onto µ-slide (80826, ibidi) at density of 2,000 cells/cm^2^ and allowed to attach overnight. Cells were then treated with SMase as previously described. Following the treatment, MSCs were stained with 5µg/mL NucBlue Live Ready Probes Reagent (Hoechst 33342, ThermoFisher) at 37℃ for 10 minutes to access nuclei. For cell imaging, differential phase contrast (DPC) images were sourced using an illumination-based DPC microscope system as described in previous studies [17]. Images were captured using a 10X objective lens followed by reconstruction. After converting the reconstructed imaging to 16-bit for processing in ImageJ, background noise was removed by using the built-in ‘background subtraction’ function in ImageJ. After filling the gaps within cells by using ‘dilation’ module in CellProfiler, a global threshold strategy with a robust background thresholding method was used to identify the cell outlines and then segment the single cells by using the customized CellProfiler pipeline. Overall, 24 morphological features were collected: Area, BoundingBoxArea, BoundingBoxMaximum_X, BoundingBoxMaximum_Y, BoundingBoxMinimum_X, BoundingBoxMinimum_Y, Center_X, Center_Y, Compactness, ConvexArea, Eccentricity, EquivalentDiamter, Extent, FormFacter, MajorAxisLength, MaxFeretDiameter, MaximumRadius, MeanRadius, MedianRadius, MinFeretDiameter, MinorAxisLength, Orientation, Perimeter, and Solidity. The features were normalized using the *StandardScaler* function in Scikit-learn 1.3.1 in Python 3.9, which standardizes each feature to zero mean and unit variance. Normalized features were then passed through unsupervised uniform manifold approximation and projection (UMAP) with optimized hyper-parameters according to the manual [41]. The first two embeddings calculated through UMAP were used for visualization of the spatial distribution in UMAP space among different groups. Component UMAP1 was used to color code the cells using Python scripts with customization.

### Secretome Analysis

Untreated (n = 3) and SMase-treated (n = 4) MSCs were cultured and treated and previously described before the conditioned media was removed and spun down at 500g to remove any debris before freezing the supernatant at -80°C. The culture media was evaluated by a custom Luminex Multiplex enzyme-linked immunosorbent assay for 39 human chemokines, cytokines, and growth factors (Thermo Fisher Scientific). Unconditioned media was used for background subtraction.

### Indolemaine2,3-Dioxygenase (IDO) Activity Assay

MSCs were seeded into a 12 well plate at a density of approximately 28,000 cells/cm^2^. The SMase group was treated with 0.3 U/mL SMase for 24 hours. SMase was removed and was washed with PBS. Next, 50 ng/mL IFN-γ was added to the positive control group (IFN-γ only) and the SMase group for 24 hours. Conditioned media was collected from each group to measure kynurenine concentration. To measure kynurenine concentration, samples were pipetted into a V-bottom well plate with a standard curve. 30% trichloroacetic acid was added to each well before spinning the plate at 950g for 5 minutes to pellet the precipitate. 75 μL of samples and standards, and 75 μL of Ehrlich’s Reagent was added to a flat bottom well plate. The plate was incubated a room temperature for 5 minutes before being read using SpectraMax M2 microplate reader (Molecular Devices, San Jose, CA) at 490 nm. Kynurenine concentration was converted to picograms of kynurenine and was normalized by cell count and number of days exposed to IFN-γ (pg kynurnine/cell/day).

### T cell Suppression Assay

T cell suppression assay was performed following a procedure described elsewhere [11]. In brief, MSCs were expanded into a T-225 flask for one passage (P2), before harvesting and seeding into a 96 well plate at a density of approximately 28,000 cells/cm^2^. The SMase group was treated with 0.3 U/mL SMase for 24 hours before being washed with PBS before the co-culture. PBMCs were labeled with carboxyfluorescein succinimydl ester according to the manufacturer’s protocol and stimulated with anti-CD3/CD28 Dynabeads (11131D, Thermo Fisher Scientifi). Approximately 10^5^ PBMCS were added to each well for a 1:1 MSC:PMBC ratio. Unstimulated PBMC control was also assessed. MSCs and PBMCs were co-cultured for 72 hours before harvesting and assessed with the Cytek Aurora.

### Macrophage Activation Assay

Macrophage activation assay was performed using a modified procedure described elsewhere [42]. THP-1s (TIB-202, ATCC, Manassas, VA) were thawed in 20% FBS, 1% P/S, 0.1% β-mercaptoethanol (21985023, Thermo Fisher Scientific) in RPMI 1640 containing L-glutamine and 10 mM HEPES (30-2001, ATCC). THP-1s were spun at 150×g for 5 min to pellet before resuspending in 12 mL of THP-1 media (about 7×10^5^ cells/mL). THP-1s were then split between two T-25 flasks and stood upright in the incubator for 24 hours. To differentiate THP-1s, 5 μL of 5 mg/mL of Phorbol 12-myristate 13 acetate (PMA) (P1585, Sigma-Aldrich) was added to 2.5 mL of RPMI 1640 media (no FBS or P/S) and filter sterilized using a syringe filter. The PMA solution was then added to media (1:100 dilution) containing 1% heat inactivated (HI) FBS (SH30071.01, Cytiva), P/S, and RPMI-1640 (differentiation media). The THP-1s were counted then harvested and spun down at 150×g for 5 minutes. The supernatant was aspirated, and the pellet was resuspending in differentiation media to give a concentration of 1.1×10^5^ THP-1s/200 μL. THP-1s are then seeded in a flat bottom 96 well plate and placed in incubator for 48 hours to generative naïve macrophages. After 48 hours, media was carefully pipetted to remove the differentiation media without disturbing the monolayer. Pro-inflammatory media was made, consisting of 25 μL IFN-γ (300-02, Thermo Fisher Scientific) (100 μg/mL stock) (1:1000 dilution) and 2.5 μL LPS (L4391, Millipore Sigma) (1 mg/mL stock) (1:10,000 dilution) was added to 25 mL of media (1% HI FBS, 1% P/S, in RPMI-1640). MSCs were harvested and added to pro-inflammatory media at 10^5^ MSCs/ 200 μL before being added to the wells containing naïve macrophages. The co-culture was gently redistributed using a multichannel pipet. The plate was then placed back in the incubator for 24 hours. After 24 hours, the well plate containing the co-culture was spun at 500×g for 5 minutes. Approximately 150 μL of conditioned media was carefully removed and placed in a V-bottom well plate and was stored in the -20C until a TNF-α ELISA (KHC3011, Invitrogen) was conducted. Briefly, 150 μL of pro-inflammatory media was added back into the wells and approximately 15 μL of conditioned media was used for the ELISA. Macrophage polarization assay was performed using a procedure described here [42]. Flow cytometry analysis was conducted using the Cytek Aurora.

### Quantification and Statistical Analysis

Statistical analysis was conducted using GraphPad Prism version 10.3.0 for Windows (GraphPad Software, Boston, Massachusetts USA, www.graphpad.com). Comparisons between groups were performed using unpaired t-tests, one-way analysis of variance with Tukey’s post-hoc pairwise tests for multiple comparisons, or two-way analysis of variance with repeated measures, where applicable. Significance was determined by p<0.05 and all data is represented as mean ± SEM on bar plots. Dimensionality reduction was performed using Python 3.9.

## RESULTS

### SMase Treatment Preserves MSC Viability and Minimally Affects Cell Death Profiles

To determine whether sphingomyelinase (SMase) treatment influences mesenchymal stem cell (MSC) survival, we first examined overall cell viability and death pathways (Figure 1). Fluorescence microscopy revealed comparable cell density and morphology between untreated (UT) and SMase-treated MSCs (Figure 1A). Quantitative assessments demonstrated no significant difference in viability (Figure 1B) or normalized proliferation (Figure 1C) between groups, indicating that exogenous SMase exposure does not diminish MSC survival. Flow cytometric analysis of Apotracker^TM^ and Zombie UV staining supported these observations; the fraction of live, early apoptotic, late apoptotic, or necrotic cells remained similar across treatments (Figure 1D–E). β-galactosidase staining showed no increase in senescent cells in SMase-treated cultures (Figure 1F–G). Moreover, autophagosome imaging and quantification revealed no significant changes in autophagy following SMase treatment (Figure 1H-J). These data collectively suggest that SMase treatment does not affect MSC viability or induce cellular and senescence.

**Figure 1.**
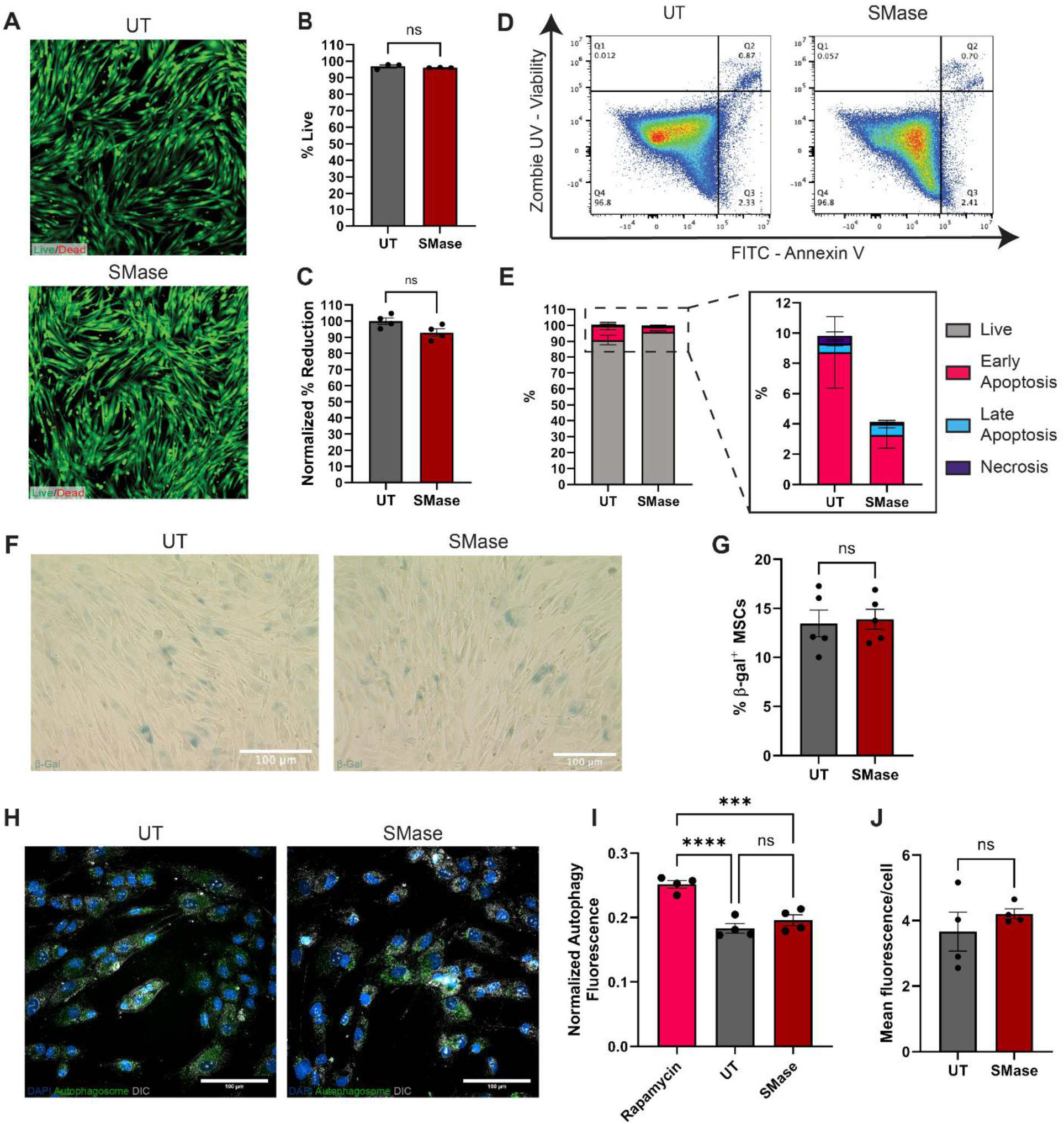
Sphingomyelinase treatment maintains MSC viability and senescence while modulating cell death profiles. (A) Representative fluorescence microscopy images of untreated (UT) and sphingomyelinase (SMase) treated MSCs stained for Live/Dead (scale bar = 100 μm). (B) Quantification of live cells after 24-hr SMase treatment between UT and SMase-treated MSCs (ns = not significant). (C) Normalized Alamar Blue percent reduction between UT and SMase-treated MSCs. (D) Representative flow cytometry plots of Annexin V/Zombie UV staining in UT and SMase-treated MSCs. (E) Quantification of cell death profiles between treatment groups. (F) Representative β-galactosidase staining images of UT and SMase-treated MSCs (scale bar = 100 μm). (G) Quantification of β-galactosidase positive cells between UT and SMase-treated MSCs (ns = not significant). (H) Representative fluorescence microscopy images showing nuclear staining (DAPI, blue), autophagosomes (green), and DIC (grey) in UT and SMase-treated MSCs (scale bars = 100 μm). (I) Fluorescence microscopy quantification of autophagosome formation following 24-hr SMase treatment between UT and SMase-treated MSCs (ns = not significant). (J) Plate reader quantification of autophagosome formation following 24-hr SMase treatment between UT and SMase-treated MSCs (ns = not significant). Data presented as mean ± SEM; statistical significance determined by unpaired t-test or one-way ANOVA.

### Mitochondrial Architecture Is Maintained Following SMase Stimulation

Given the regulatory role ceramides play in mitochondrial function, we investigated whether exogenous SMase affects mitochondrial morphology and network structure (Figure 2). Filamentous mitochondria with interconnected networks were observed in both UT and SMase-treated MSCs from confocal imaging (Figure 2A). Quantitative 3D analyses revealed no significant differences in mean mitochondrial volume and sphericity between the two groups (Figure 2B-C), indicating that SMase treatment did not alter overall mitochondrial morphology. Analysis of mitochondrial networks showed the number of branches per mitochondrion were comparable between the groups (Figure 2D). Although the treated cells exhibited a significant increase in mean branch length (Figure 2F), this did not result in a difference in total branch length (Figure 2E). Additionally, there was a trend toward increased mean branch diameter in the treated group (Figure 2G, p=0.0630), which is consistent with the modest, non-significant increase in mean mitochondrial volume (Figure 2B). These mitochondrial insights suggest lipid alterations mediated through exogenous SMase delivery do not cause apparent morphological differences in mitochondria.

**Figure 2.**
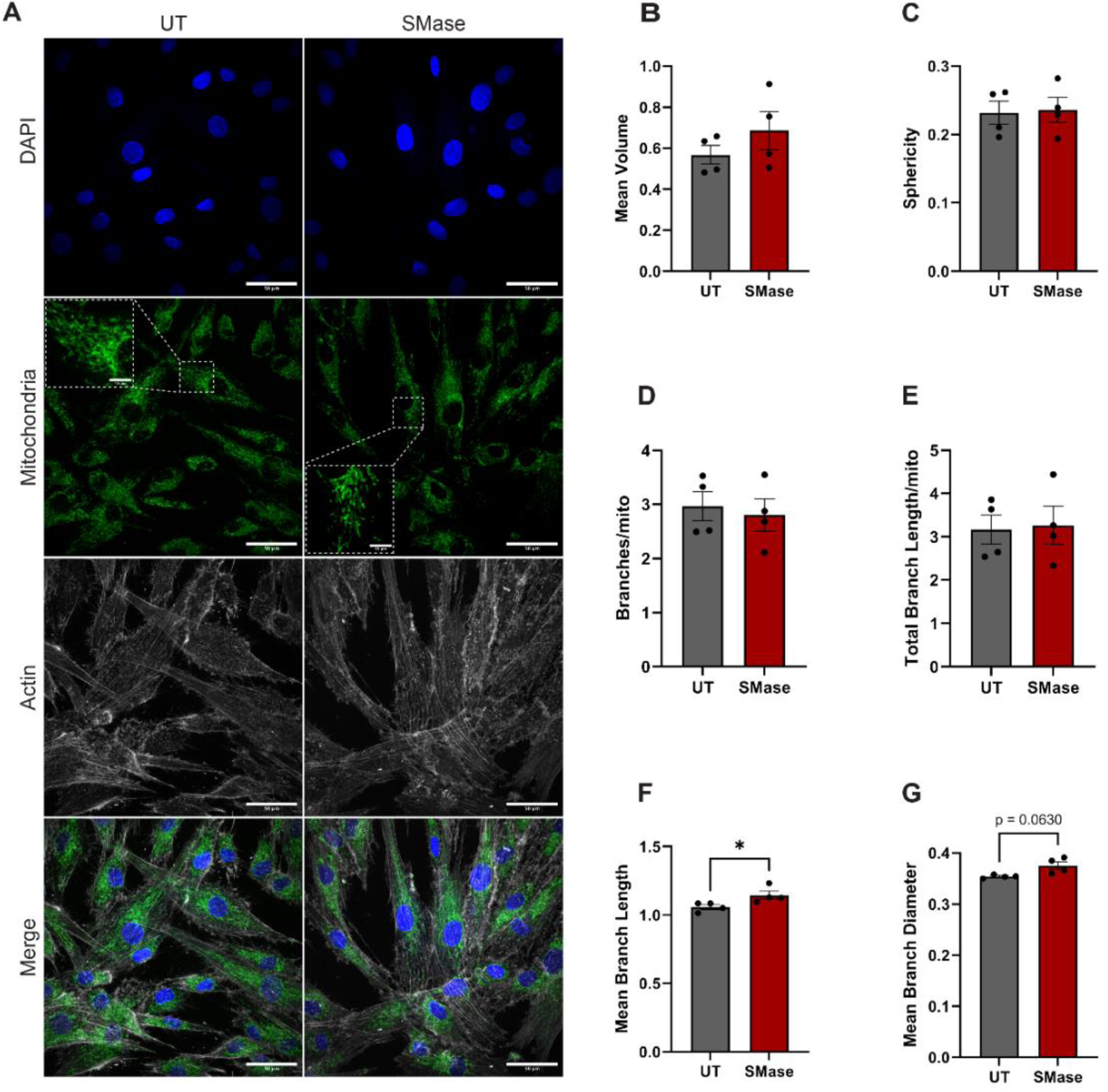
MSC mitochondrial networks and morphology characteristics are intact following SMase treatment. (A) Representative fluorescence microscopy images showing nuclear staining (DAPI, blue), mitochondrial networks (green), and F-actin cytoskeleton (gray) in untreated (UT) and SMase-treated MSCs (scale bars = 50 μm). Insets show magnified regions of mitochondrial morphology (scale bar = 10 μm). (B) Quantification of mean mitochondrial volume between treatment groups. (C) Analysis of mitochondrial sphericity comparing UT and SMase-treated cells. (D-E) Evaluation of mitochondrial branch characteristics including (D) branches per mitochondrion ratio and (E) total branch length per mitochondrion. (F) Measurement of significant mean branch length in mitochondrial networks. (G) Quantification of mean branch diameter between UT and SMase treatment. Data presented as mean ± SEM; statistical significance determined by unpaired t-test; (*) *p* < 0.05.

### Reprograming of Sphingolipid Metabolism Occurs following SMase Treatment

To explore the metabolic effects of exogenous SMase stimulation on several key lipids, we conducted a time series metabolic tracer workflow to image the persistence of fluorescently labeled NDB-lipids and quantified these metabolized lipids through HPLC analysis (Figure 3A). Preliminary experiments found that following C6 NBD-SM incubation, strong green fluorescence was observed in both groups initially, however over time, there was a quicker decrease of fluorescence in UT MSCs shown at 3 hours, compared to SMase pre-treated MSCs, shown by the lack of signal by the 24-hour timepoint (Figure 3B). We collected the samples at each timepoint (0, 1, 3, 5, 9, and 24-hours) and conducted lipid extraction and total protein analysis for normalization. HPLC analysis was conducted for all samples to identify NBD-fatty acid (NBD-FA), NBD-sphingomyelin (NBD-SM), NBD-ceramide (NBD-Cer), and NBD-glucosylceramide (NBD-GlcCer) concentrations (Figure 3C). We found there weren’t any differences in NBD-FA levels between UT and SMase-treated MSCs. However, both groups saw an overall decreasing NBD-FA trend over time. When assessing NBD-SM in cells, at timepoint 0 hours, UT MSCs seemed to have higher levels of NBD-SM compared to SMase-treated MSCs. Interestingly, between timepoints 1-24 hours SMase-treated MSCs had higher levels of NBD-SM compared to UT MSCs with statistical significance being shown at the 1- and 5-hour marks (p= <0.05 and p=<0.01, respectively). Cellular NBD-Cer levels were generally comparable between both groups. However, SMase-treated MSCs had a greater concentration at 1 and 5 hours (p= <0.05). When monitoring NBD-GlcCer levels in cells, we saw higher levels (non-significant) of NBD-GlcCer in SMase-treated MSCs suggesting that intracellular NBD-Cer is converted to NBD-GlcCer. Looking at the overall lipid composition of the main 3 species (SM, Cer, and GlcCer), we see at time 0, there were similar levels of SLs between both groups. However, there were lower levels of measured intracellular NBD-lipids in UT MSCs compared to SMase-treated MSCs over time (Figure 3D).

**Figure 3.**
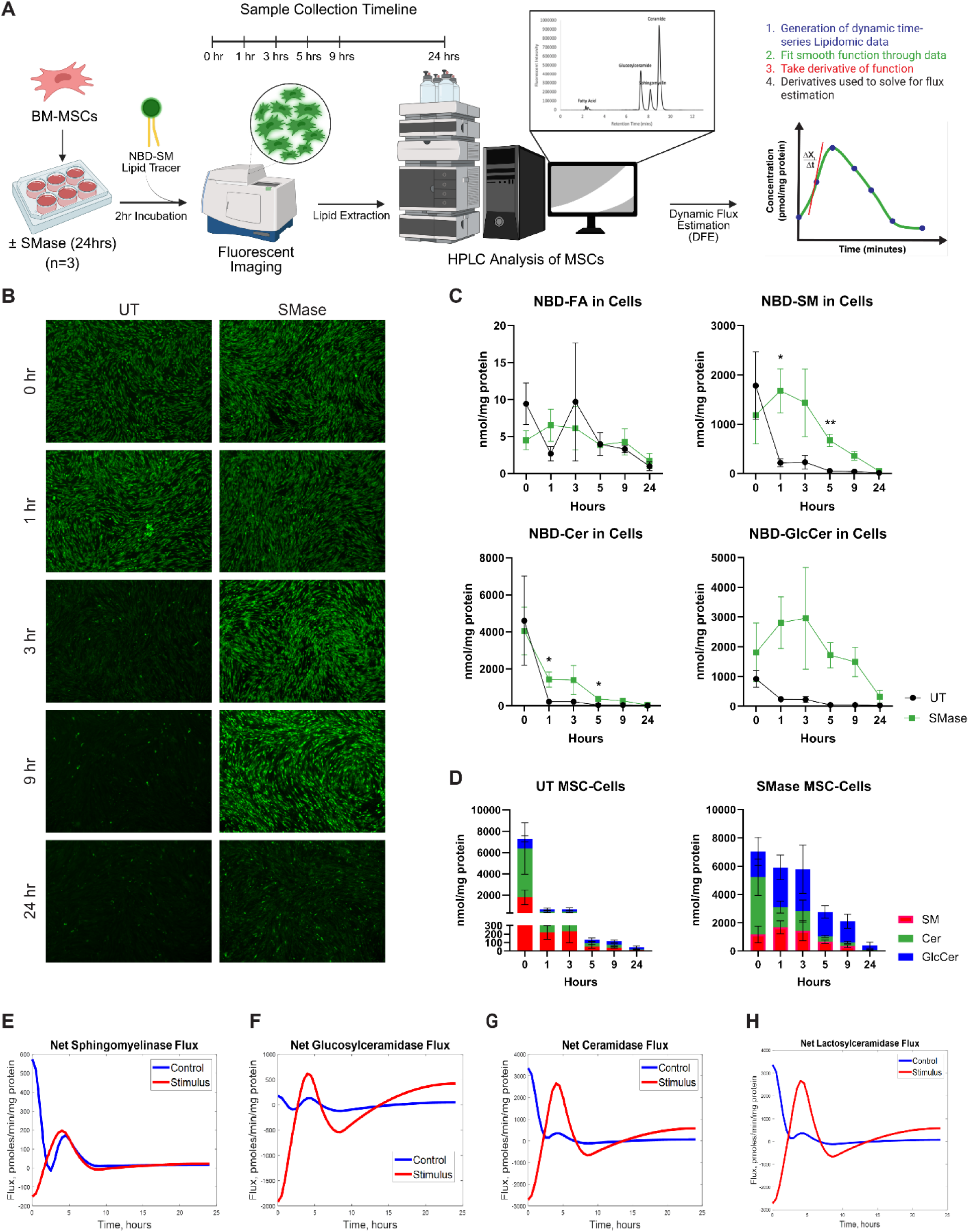
MSC metabolism of NBD-labeled lipids is altered following SMase treatment. (A) Experimental overview of NBD-tracer studies, mathematical workflow, and principles involved in the DFE model. (B) Fluorescent microscopy images of NBD-tracer over time in UT and SMase treated MSCs. (C) Quantification of lipid concentration over time within BM-MSCs following NBD-SM addition in UT and SMase treated MSCs. (D) Stacked bar plots of quantified NBD-labeled lipids in MSCs. Net enzymatic flux in response to 24-hour SMase simulation for (E) Sphingomyelinase, (F) Glucosylceramidase, (G) Ceramidase, and (H) Lactosylceramidase. Data presented as mean ± SEM. Statistical significance determined by unpaired t-test with Welch’s correction where appropriate; *p < 0.05, **p < 0.01.

Using a Dynamic Flux Estimation (DFE) model we were able to predict the net metabolic fluxes of SMase, glucosylceramidase (GLUCDase), ceramidase (CDase), and lactosylceramidase (LACCDase) using dynamic timeseries lipid concentrations described previously. When we look at the overall net SMase flux model where a negative flux along the y-axis represents an opposing reaction (e.g. negative flux = sphingomyelin synthase (SMS) activity dominates, positive flux = SMase activity dominates), we initially see SMase activity dominating the fluxes in UT MSC while SMS activity dominates the SMase pre-treated MSCs, with SMase activity persistent during the remainder of the time (Figure 3E). Estimation of the net GLUCDase flux (positive flux = GLUCDase activity dominates, and negative flux = glucosylceramide synthase (GLUCS) activity dominates) shows initially GLUCS dominating the enzymatic activity in SMase-treated MSCs and GLUCDase activity dominating the UT group (Figure 3F). However, during the 3-to-9-hour timepoints, net activity fluctuates ultimately leading to higher GLUCDase activity in the SMase stimulated group at 24 hours. When estimating the net CDase flux (Figure 3G) we see similar trends with initial positive activity (CDase-driven) in the UT group and ceramide synthase (CERS-driven) activity driving the reaction in the SMase-treated group. Approaching 24 hours the estimate shows CDase to be active in the SMase-treated group. Lastly, LCS activity is initially apparent in SMase group with the opposite enzyme lactosylceramidase synthase (LACCDase) active in the UT group. Similarly, the enzymatic reactions flip throughout the 3-to-9-hour time period between LCS and LACCDase with LACCDase leading the activity by the 24-hour mark in the SMase-treated group. Overall, the SMase-treated MSC group has a net hydrolytic flux pattern at 24 hours with LACCDase, GLUCDase, and CDase all dominating.

### Morphological Screening Points to an Immunomodulatory-like State following SMase Treatment

To assess whether SMase treatment induces morphological changes consistent with an immunomodulatory phenotype, we performed high resolution imaging and quantitative morphometric analysis of MSCs (UT, SMase-treated, or IFN-γ-treated). As illustrated in Figure 4A and previously shown [43], a total of 24 morphological features were extracted from segmented single cells through CellProfiler and compared across treatment conditions, including cell area, perimeter, feret diameters, form factor, compactness, solidity, etc. Unsupervised dimensionality reduction strategy uniform manifold approximation and projection (UMAP) was used to display MSC morphological variation and reveal distinct clustering of MSCs by treatment condition (Figure 4B). All groups showed independent clustering with UT cells on the left of UMAP1, IFN-γ–treated cells positioned on the right of UMAP1, and SMase-treated cells clustered between the two. The left-to-right shifting in morphology suggests that MSCs respond to SMase priming in a manner partially similar to the response to IFN-*γ*. To further explore these morphological signatures, multiple key individual features were overlaid on the UMAP (Figure 4C). Notably, SMase-treated cells displayed high values similar to IFN-γ–treated cells in area, perimeter, max ferret diameter, and compactness relative to UT groups, suggesting an increase in cell size and elongation with priming. Representative color-coded DPC images corresponding to each treatment condition, overlaid with UMAP1 (Figure 4D), illustrate visual concordance between cellular phenotype and computational clustering.

**Figure 4.**
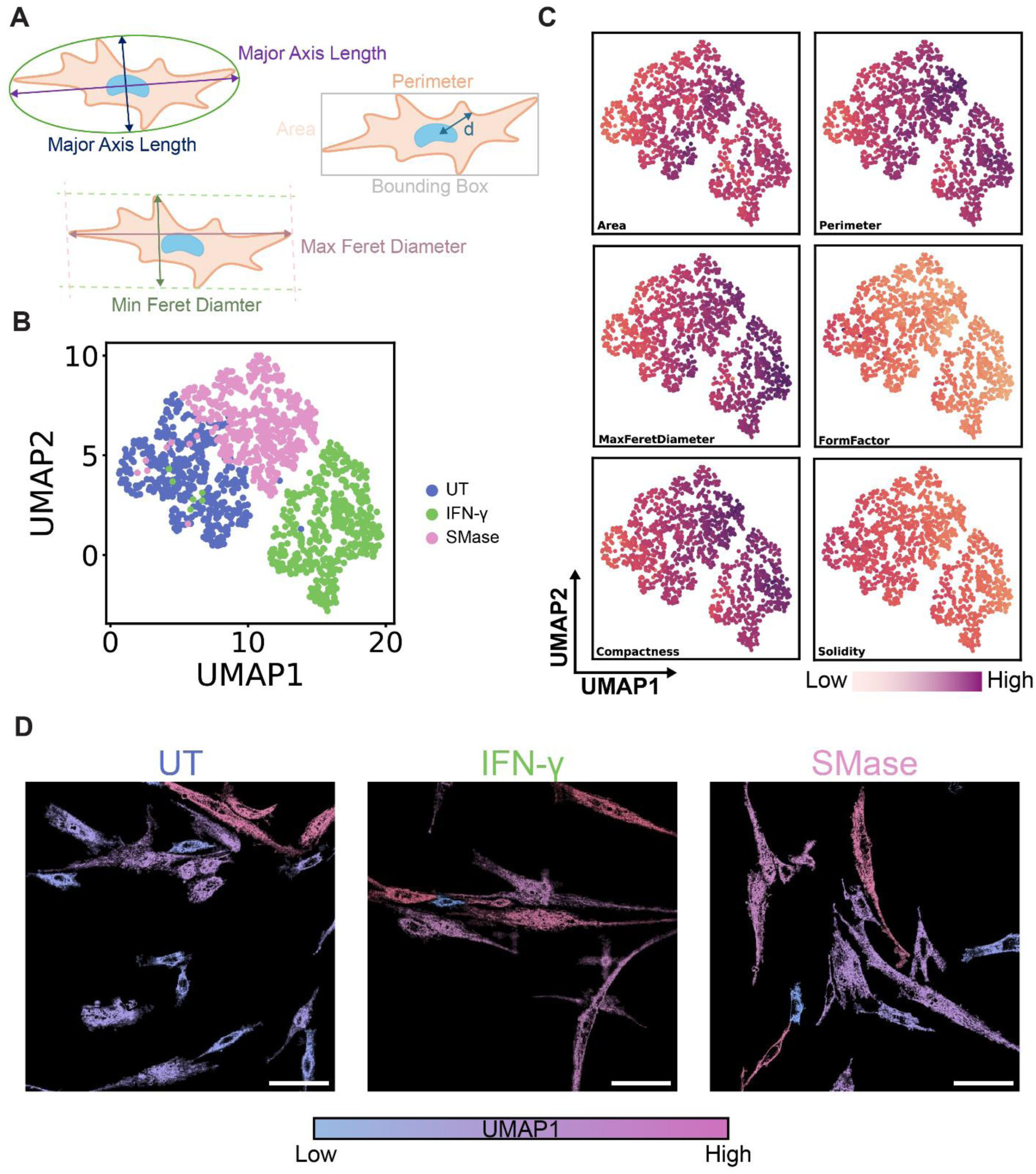
Morphological assessment of MSCs uncover SMase treatment sways morphology characteristics toward an immunomodulatory-like state. (A) Morphological features measured in cell profiler. (B) UMAP of morphology features amongst UT, IFN-γ, and SMase treated cells. (C) UMAP of parameters computed using measured morphological features. (D) Fluorescent microscopy images of MSCs (scale bars = 100 μm) overlaid with UMAP1 (blue = low values, magenta = high values).

### SMase Treatment Induces Distinct Shifts in the MSC Secretome

Given the importance of MSC paracrine function, we profiled the secretome in UT vs. SMase-treated cells (Figure 5). Hierarchical clustering of soluble factors revealed two prominent clusters that distinguished UT and SMase samples (Figure 5A). Multiple immunomodulatory cytokines (e.g., IL-4, IL-7, IL-10) and chemokines (SDF-1α, IP-10, RANTES) were significantly downregulated upon SMase treatment (Figure 5B), accompanied by reduced levels of MMP-2 and PDGF-BB. In contrast, expression of factors such as IL-15, TNF-α, GM-CSF, and MIP-1α remained unchanged between groups (Figure 5C). Strikingly, proangiogenic mediators (VEGF-A, VEGF-D, HGF, EGF) and certain inflammatory markers (IL-6, IL-8, Eotaxin, M-CSF) were significantly upregulated under SMase treatment (Figure 5D). These data indicate that exogenous SMase reconfigures the MSC secretome toward enhanced growth factor and chemokine release, potentially modulating extracellular vesicle content and the local microenvironment.

**Figure 5.**
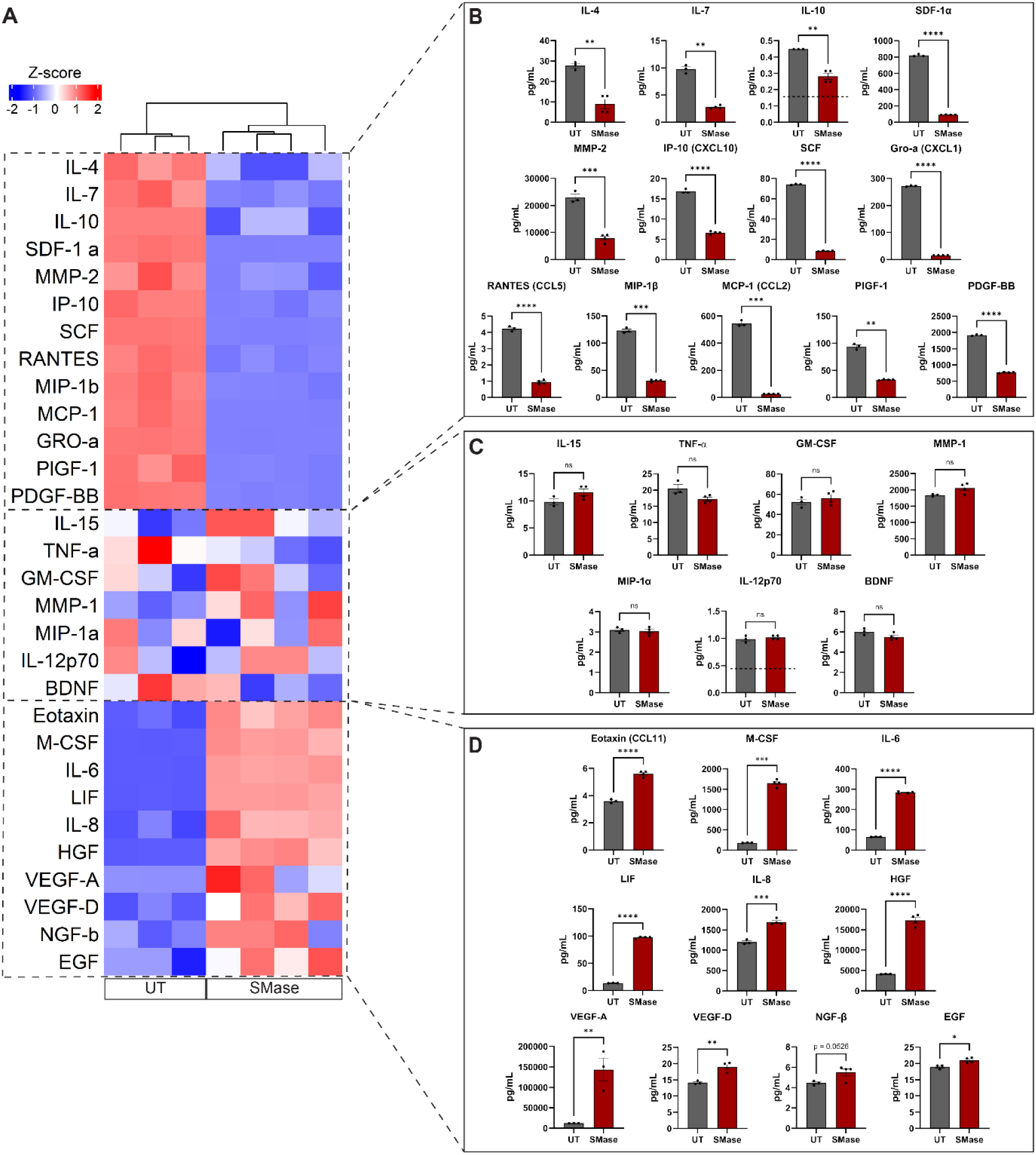
SMase treatment significantly alters MSC secretome profile with distinct modulation of pro-inflammatory and growth factors. (A) Hierarchical clustering heatmap showing differential expression of secreted factors between untreated (UT) and SMase-treated MSCs. Color scale represents normalized expression levels (blue: decreased; red: increased). (B) Quantification of significantly downregulated factors in SMase-treated MSCs, including key immunomodulatory cytokines (IL-4, IL-7, IL-10), chemokines (SDF-1α, IP-10, RANTES), and growth factors (MMP-2, PDGF-BB). (C) Analysis of factors showing no significant changes between treatment groups, including IL-15, TNF-α, GM-CSF, MMP-1, MIP-1α, IL-12p70, and BDNF. (D) Quantification of significantly upregulated factors in SMase-treated MSCs, highlighting increased expression of pro-angiogenic and growth factors (VEGF-A, VEGF-D, HGF, EGF), inflammatory mediators (IL-6, IL-8), and chemokines (Eotaxin, M-CSF, LIF). Data presented as mean ± SEM. Statistical significance determined by unpaired t-test; *p < 0.05, **p < 0.01, ***p < 0.001, ****p < 0.0001.

### MSCs Retain Immunomodulatory Functions Following SMase-Mediated Lipid Alterations

To assess if SMase has an effect of MSC immunomodulatory potential, we conducted T cell proliferation, IDO activity, and macrophage assays (Figure 6A). An assessment of IDO activity was conducted by measuring kynurenine (kyn) levels in conditioned media, a product of tryptophan metabolism via IDO. SMase treated MSCs produced 17.40 ± 1.40 pg kyn/cell/day, slightly less than the IFN-γ positive control (18.29 ± 1.68 vs. 17.40 ± 1.40, ns). Both the IFN-γ- and SMase-treated cells had greater IDO activity compared to the unstimulated, negative control group (0.36 ± 0.05 pg kyn/cell/day, p < 0.001) (Figure 6B). Next, we evaluated the ability of treated MSCs to reduce T-cell proliferation in vitro. After 72 hours of co-culture with peripheral blood mononuclear cells, there was 28.86 ± 2.06% CD4 T cell proliferation with UT MSCs (p < 0.0001). With SMase-treated MSCs, there was 44.42 ± 2.54% CD4 T proliferation (p < 0.0001). There was a statistically significant difference in CD4 T cell suppression between UT and SMase treated MSCs (p < 0.01) (Figure 6C). CD8 T cells had 34.64 ± 2.92% proliferation after co-culture with UT MSCs while SMase-treated MSCs there was 58.78 ± 1.47% proliferation. There was statistically significant difference in CD8 T cell suppression between UT and SMase, where SMase-treated MSCs had less suppression compared to UT MSCs (p < 0.0001) (Figure 5D).

**Figure 6.**
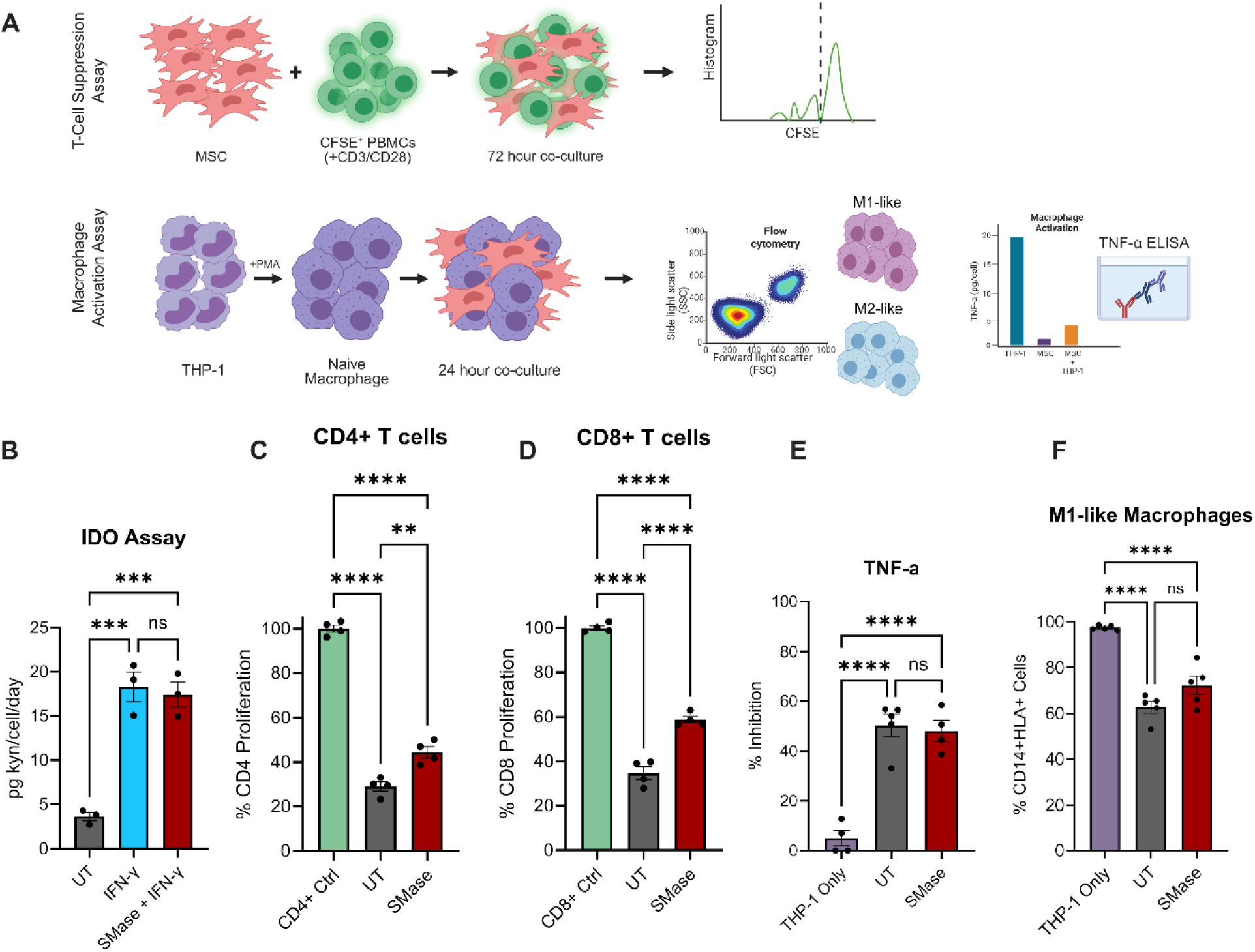
SMase treated MSCs lead to reduced immune cell activation in T-cells and Macrophages. (A) Schematic representation of experimental workflow. (B) Quantification of kynurenine from IDO activity assay. (C) Percent proliferation of CD4+ T cells after co-culture with MSCs. (C) Percent proliferation of CD8+ T cells after co-culture with MSCs. (D) Percent inhibition of TNF-α secretion from THP-1 derived macrophages after 24-hour co-culture with MSCs. (E) Percent CD14+HLA-DR+ THP-1s after 72-hour co-culture with MSCs. (A) n = 4, (B) n = 4, (C) n = 3, (D) n = 4 (E) n = 4. Data presented as mean ± SEM. Statistical significance determined by one-way ANOVA with multiple comparisons test; *p < 0.05, **p < 0.01, ***p < 0.001, ****p < 0.0001.

We assessed macrophage activation and polarization by measuring TNF-α inhibition and surface marker expression, respectively. After 24-hour MSC: THP-1 derived macrophage co-culture, UT MSCs had 50.23% ± 4.42% TNF-α inhibition. SMase-treated MSCs had 48.06% ± 4.23% TNF-α inhibition. When comparing TNF-α inhibition to the THP-1 macrophage only control, both groups statistically significantly inhibited TNF-α levels (p < 0.0001) (Figure 6E). There was no statistically significant difference between UT and SMase-treated MSCs. Next, we assessed macrophage polarization after 72-hour MSC: THP-1 co-culture by measuring percent CD14+HLA-DR+ cells, representing M1-like macrophages. UT MSCs had a lower percentage of CD14+HLA-DR+ cells compared to THP-1 only control (62.72 ± 2.60% vs. 97.42 ± 0.46%, p < 0.0001). Additionally, SMase-treated MSCs had a lower percentage of CD14+HLA-DR+ cells compared to THP-1 only control (72.22 ± 3.98% vs. 97.42 ± 0.46%, p < 0.0001). There was no statistically significant difference between UT and SMase-treated MSCs (Figure 6F).

## DISCUSSION

MSCs remain among the most widely researched cell types for regenerative medicine and immunotherapy due to their capacity for immunomodulation, secretion of trophic factors, and support of tissue repair. Despite their biological promise, the clinical translation of MSCs has been hindered by inconsistent potency and manufacturing variability. One of the most critical challenges in advancing MSC therapy is the functional heterogeneity of MSC populations. Donor variability, culture conditions, and lack of standardized potency assays all contribute to inconsistent clinical outcomes. To overcome this, various priming strategies have been investigated to “license” MSCs temporarily or stably inducing them into a therapeutically active state. Traditional cytokine-based priming approaches using IFN-γ, TNF-α, or interleukin-1β have been shown to upregulate key immunosuppressive mediators such as indoleamine 2,3-dioxygenase (IDO), prostaglandin E2 (PGE2), and PD-L1, thereby enhancing their anti-inflammatory effects in preclinical models of GVHD, colitis, and other immune-mediated conditions. In this work, we aim to identify the use of lipid metabolic perturbation as a means to alter MSC phenotype and support their immunomodulatory capabilities.

Our data indicate treating MSCs with exogenous SMase preserves viability while selectively modifying cellular architecture. The negligible impact on both short- and long-term cell survival indicates that SMase activity does not adversely affect MSC viability or senescence profiles, thereby supporting the feasibility of enzymatic manipulation for therapeutic use. In a study conducted by Ogle et al., an assessment of hydrogel stiffness on MSC secretome and senescence was conducted. When evaluating MSC senescence, they observed P2 MSCs grown on tissue culture plastic had approximately 15% β-gal+ senescent cells, similar to our results [44]. These findings align with previous studies showing that sphingolipid perturbations can modify cellular behavior without compromising viability [25, 45]. Interestingly, the preservation of elongated and interconnected mitochondrial morphology, reflected by consistent volume, sphericity, and network parameters, implies that SMase treatment does not induce stress or disrupt the mitochondrial dynamics essential for meeting cellular energy demands [46]. The observed increase in mean branch length and diameter, without changes to total branch length and volume, may indicate a subtle compensatory mechanism that helps sustain mitochondrial connectivity and function. Further exploration through experimentation to elucidate the functional implications of these structural findings, such as mitochondrial bioenergetics, could provide deeper insights into the impact of SMase on MSC mitochondrial physiology.

Often, we think of lipid metabolism as a steady process within a cell where a lipid is converted from one species to another, resulting in a cell response. In actuality, it is a dynamic process of intracellular lipid metabolism with multiple lipid influxes and effluxes. This phenomenon emphasizes the importance of culture conditions and lipid metabolites in media formulations as these lipids can influence intracellular lipid metabolism and overall cell behavior. SMase-treated MSCs exhibit sustained metabolic reprogramming up to 36–48 hours post-treatment, as indicated by integrated lipidomic and dynamic flux estimation (DFE) modeling. Unlike transient responses often observed in cytokine-stimulated cells, SMase induces broader and more robust changes in the lipid metabolic state. To build upon our findings from this work, we aim to understand how SL fluxes influence MSC behavior while taking variables such as culture conditions, donor variability, and tissue source into consideration. Using flux balance equations, we developed a preliminary DFE model. DFE predictions revealed persistently altered metabolic net flux distributions in GLUCDase, CDase, and LACCDase, patterns not observed in untreated cells. In a study by Chiappa et al., a DFE model was utilized to uncover putative SL targets (SMS, Sphingosine Kinase, and CDase) for the resolution of inflammation following KDO_2_-Lipid A (a key component of lipopolysaccharide found in most gram-negative bacteria) stimulated macrophages [47]. In addition to serving as targets, these SL enzymes have been shown to be central to cellular functions such as differentiation and vascular remodeling. Yang et al. identified the integral role of SMS in the facilitation of MSC osteogenic differentiation and promotion of angiogenesis through modulating the Cer/PP2A/Akt pathway [48]. Alongside SL enzymes, SLs such as glycosphingolipids, C1P, sphingosine (Sph) and S1P play various roles in the transcription of genes, chromatin accessibility and remodeling, and epigenetic modifications [49, 50]. For example, Sph can bind to steroidogenic factor-1 (SF-1) and co-repressor protein complexes, preventing the transcription of target genes [51]. Additionally, SMase may influence transcription indirectly through the production of Cer, decreasing DG levels. For example, DG is converted to phosphatidic acid via diacylglycerol kinase activity. PA then can bind to SF-1, activating its receptor and the recruitment of co-activator proteins, resulting in transcription of its target genes [52]. Conducting lipidomics and techniques such as Assay for Transposase-Accessible Chromatin (ATAC)-seq can provide insight into if Cer and its downstream metabolites are affecting chromatin structure and transcription.

In addition to lipid metabolic reprogramming, we observed pronounced changes in the MSC secretome following SMase treatment. The altered profile includes several cytokines and soluble factors associated with both immune suppression (IL-4 and IL-10) and tissue repair (VEG-A and VEGF-D). This mirrors prior findings showing that priming can modulate not only intracellular pathways but also the composition of extracellular vesicles and soluble proteins, which are key effectors of MSC therapeutic action. MSCs can modulate effector T cell and macrophage behavior via IDO activity, cell-cell contact mechanisms, or through paracrine factors. We saw less T cell suppression from SMase-treated MSCs compared to UT MSCs. This may have been caused by decreased secretion of anti-inflammatory factors (IL-4, IL-10, MCP-1) and increased IL-6 secretion. We observed no changes in MSC IDO activity, suggesting that the decrease of T cell suppression was likely due to other biological mechanisms. When assessing macrophage activation, both conditions inhibited TNF-α after 24-hours and decreased the percentage of M1-like macrophages after 72-hour MSC: THP-1 co-culture.

An additional dimension of our analysis involved quantifying morphological features of MSCs as potential proxies for functional state. High-content imaging and UMAP embedding of morphological features revealed that SMase-treated MSCs adopt a cell shape profile partially overlapping with IFN-γ–primed cells. These cells exhibited decreased solidity and FormFactor, increased compactness and max feret diameter, indicating elongation and irregularities in the cell shape. Prior research by Larey et al. has established correlations between MSC morphology and immunomodulatory potency, where more compact, less spread morphologies are associated with anti-inflammatory function [53]. Additionally, Priyadarshani et al. uncovered distinct changes in MSC function were linked to higher max ferret diameter, compactness, and perimeter. These same changes were also parallel to shifts in the lipidome with higher levels of phosphatidylcholines, lysophosphatidylcholines, and triacylglycerol [17]. Taken together, our findings reinforce the idea that morphological metrics can serve as predictive, non-destructive quality attributes for MSC potency following exogenous licensing.

### Limitations of the study

While this study identifies SMase as a novel priming agent capable of safely inducing changes in morphological, metabolic, and functional changes in MSCs, several limitations should be considered. SMase represents only one of several enzymes involved in sphingolipid metabolism that can influence ceramide generation; future studies should explore the effects of other ceramide-producing enzymes such as ceramide synthases or glucosylceramidase to determine whether distinct pathways yield similar or complementary functional states. In addition, SMase exposure in this study was limited to a single 24-hour treatment window. The long-term persistence of the primed phenotype and the potential benefits of extended or cyclic SMase stimulation remain unexplored and warrant systematic investigation. All experiments were performed using MSCs derived from a single donor and tissue source. Given the known variability in MSC phenotype and function across donors and anatomical origins, additional studies incorporating multiple donors and sources are essential to assess the priming effect of SMase. Lastly, while our results indicate mitochondrial morphology is unaltered following SMase treatment, the precise impact on mitochondrial function remains undefined. Dedicated bioenergetic profiling such as mitochondrial respiration and ATP production assays will be necessary to fully characterize the influence of SMase priming on MSC mitochondrial health and metabolic fitness.

## ACKNOWLEDGEMENTS

Funding was provided by the National Science Foundation Engineering Research Center on Cell Manufacturing Technologies (EEC1648035). S.A.D. and D.C.S. were supported by the NGMS-sponsored Cell and Tissue Engineering NIH Biotechnology Training Grant (T32 GM-008433 and T32 GM145735, respectively). The authors would like to thank the Parker H. Petit Institute for Bioengineering and Biosciences at Georgia Institute of Technology for shared equipment and core services. Figures were created using BioRender.com and Adobe Illustrator.

## AUTHOR CONTRIBUTIONS

S.A.D. and D.C.S. designed and conducted the experiments, analyzed the data, generated the figures, wrote, and revised the manuscript. K.R., H.Z., and N.L. performed experiments, analyzed data, wrote, and revised the manuscript. N.F.C and R.O. analyzed data, wrote, and revised the manuscript. E.A.B. supervised the project, generated figures, wrote, and revised the manuscript. Revising and approving the final version of manuscript: All authors. All authors have read the journal’s authorship agreement and declare no conflict of interest.

## Declaration of Interests

The authors declare no competing interests.

## Notes

### Competing Interest Statement

The authors have declared no competing interest.

## REFERENCES

1. Gao, Q., et al., Bone marrow mesenchymal stromal cells: identification, classification, and differentiation. Frontiers in cell and developmental biology, 2022. 9: p. 787118.

2. Jimenez-Puerta, G.J., et al., Role of mesenchymal stromal cells as therapeutic agents: potential mechanisms of action and implications in their clinical use. Journal of clinical medicine, 2020. 9(2): p. 445.

3. Song, N., M. Scholtemeijer, and K. Shah, Mesenchymal Stem Cell Immunomodulation: Mechanisms and Therapeutic Potential. Trends Pharmacol Sci, 2020. 41(9): p. 653–664.

4. Maacha, S., et al., Paracrine mechanisms of mesenchymal stromal cells in angiogenesis. Stem cells international, 2020. 2020(1): p. 4356359.

5. Bazzoni, R., et al., Extracellular vesicle-dependent communication between mesenchymal stromal cells and immune effector cells. Frontiers in cell and developmental biology, 2020. 8: p. 596079.

6. Wu, X., et al., Mesenchymal stromal cell therapies: immunomodulatory properties and clinical progress. Stem cell research & therapy, 2020. 11(1): p. 345.

7. Zhou, T., et al., Challenges and advances in clinical applications of mesenchymal stromal cells. Journal of Hematology & Oncology, 2021. 14(1): p. 24.

8. Mastrolia, I., et al., Challenges in Clinical Development of Mesenchymal Stromal/Stem Cells: Concise Review. Stem Cells Transl Med, 2019. 8(11): p. 1135–1148.

9. Le Blanc, K., et al., ISCT MSC Committee Statement on the US FDA Approval of Allogenic Bone-Marrow Mesenchymal Stromal Cells. 2025, Elsevier.

10. Chinnadurai, R., et al., Potency analysis of mesenchymal stromal cells using a combinatorial assay matrix approach. Cell reports, 2018. 22(9): p. 2504–2517.

11. Maughon, T.S., et al., Metabolomics and cytokine profiling of mesenchymal stromal cells identify markers predictive of T-cell suppression. Cytotherapy, 2022. 24(2): p. 137–148.

12. Kota, D.J., et al., Prostaglandin E2 indicates therapeutic efficacy of mesenchymal stem cells in experimental traumatic brain injury. Stem cells, 2017. 35(5): p. 1416–1430.

13. Campos, J., et al., Lipid priming of adipose mesenchymal stromal cells with docosahexaenoic acid: impact on cell differentiation, senescence and the secretome neuroregulatory profile. Tissue Engineering and Regenerative Medicine, 2025. 22(1): p. 113–128.

14. S’Dravious, A.D., et al., Characterizing human mesenchymal stromal cells’ immune-modulatory potency using targeted lipidomic profiling of sphingolipids. Cytotherapy, 2022. 24(6): p. 608–618.

15. Nincheri, P., et al., Sphingosine 1-phosphate induces differentiation of adipose tissue-derived mesenchymal stem cells towards smooth muscle cells. Cellular and molecular life sciences, 2009. 66: p. 1741–1754.

16. Bitterlich, L., et al., The free fatty acid palmitate enhances MSC suppression of macrophages in a ceramide/CCL2/IL-10 dependent manner. Cytotherapy, 2025. 27(5): p. S52–S53.

17. Priyadarshani, P., et al., Investigation of MSC potency metrics via integration of imaging modalities with lipidomic characterization. Cell Reports, 2024. 43(8).

18. Chatgilialoglu, A., et al., Restored in vivo-like membrane lipidomics positively influence in vitro features of cultured mesenchymal stromal/stem cells derived from human placenta. Stem Cell Research & Therapy, 2017. 8(1): p. 31.

19. Kim, C., et al., Ceramide-1-phosphate regulates migration of multipotent stromal cells and endothelial progenitor cells—implications for tissue regeneration. Stem Cells, 2013. 31(3): p. 500–510.

20. Price, S.T., et al., Sphingosine 1-phosphate receptor 2 regulates the migration, proliferation, and differentiation of mesenchymal stem cells. International journal of stem cell research and therapy, 2015. 2(2): p. 014.

21. Rajesh, M., A. Kolmakova, and S. Chatterjee, Novel Role of Lactosylceramide in Vascular Endothelial Growth Factor–Mediated Angiogenesis in Human Endothelial Cells. Circulation research, 2005. 97(8): p. 796–804.

22. Yoshida, H., et al., B4GALNT1 induces angiogenesis, anchorage independence growth and motility, and promotes tumorigenesis in melanoma by induction of ganglioside GM2/GD2. Scientific reports, 2020. 10(1): p. 1199.

23. Lee, M., S.Y. Lee, and Y.-S. Bae, Functional roles of sphingolipids in immunity and their implication in disease. Experimental & molecular medicine, 2023. 55(6): p. 1110–1130.

24. Kang, H., et al., The therapeutic effects of human mesenchymal stem cells primed with sphingosine-1 phosphate on pulmonary artery hypertension. Stem Cells and Development, 2015. 24(14): p. 1658–1671.

25. Fiorani, F., et al., Ceramide releases exosomes with a specific miRNA signature for cell differentiation. Scientific Reports, 2023. 13(1): p. 10993.

26. Wong, M.-L., et al., Acute systemic inflammation up-regulates secretory sphingomyelinase in vivo: a possible link between inflammatory cytokines and atherogenesis. Proceedings of the National Academy of Sciences, 2000. 97(15): p. 8681–8686.

27. Maceyka, M. and S. Spiegel, Sphingolipid metabolites in inflammatory disease. Nature, 2014. 510(7503): p. 58–67.

28. Tabas, I., Secretory sphingomyelinase. Chemistry and physics of lipids, 1999. 102(1-2): p. 123–130.

29. Andrews, S., et al., Priming of MSCs with inflammation-relevant signals affects extracellular vesicle biogenesis, surface markers, and modulation of T cell subsets. Journal of Immunology and Regenerative Medicine, 2021. 13: p. 100036.

30. Noronha, N.d.C., et al., Priming approaches to improve the efficacy of mesenchymal stromal cell-based therapies. Stem cell research & therapy, 2019. 10: p. 1–21.

31. Hackel, A., et al., Immunological priming of mesenchymal stromal/stem cells and their extracellular vesicles augments their therapeutic benefits in experimental graft-versus-host disease via engagement of PD-1 ligands. Frontiers in Immunology, 2023. Volume 14 **-**2023.

32. Leventhal, A.R., et al., Acid sphingomyelinase-deficient macrophages have defective cholesterol trafficking and efflux. Journal of Biological Chemistry, 2001. 276(48): p. 44976–44983.

33. Bollinger, C.R., V. Teichgräber, and E. Gulbins, Ceramide-enriched membrane domains. Biochimica et Biophysica Acta (BBA)-Molecular Cell Research, 2005. 1746(3): p. 284–294.

34. Schmieder, S., R. Tatituri, and W.I. Lencer, Synthesis of a GM1 structural library reveals distinct membrane behavior based on ceramide structure. Biophysical Journal, 2024. 123(3): p. 505a.

35. Arumugam, S., et al., Ceramide structure dictates glycosphingolipid nanodomain assembly and function. Nature communications, 2021. 12(1): p. 3675.

36. Quadri, Z. and E. Bieberich, Staying sane in the membrane: Neutral sphingomyelinase 2 as a master regulator of plasma membrane ceramide. Journal of Lipid Research, 2025. 66(2): p. 100737.

37. Schneider, C.A., W.S. Rasband, and K.W. Eliceiri, NIH Image to ImageJ: 25 years of image analysis. Nature Methods, 2012. 9(7): p. 671–675.

38. Schindelin, J., et al., Fiji: an open-source platform for biological-image analysis. Nature Methods, 2012. 9(7): p. 676–682.

39. Chaudhry, A., R. Shi, and D.S. Luciani, A pipeline for multidimensional confocal analysis of mitochondrial morphology, function, and dynamics in pancreatic β-cells. American Journal of Physiology-Endocrinology and Metabolism, 2020. 318(2): p. E87–E101.

40. Nikolova-Karakashian, M., et al., Bimodal regulation of ceramidase by interleukin-1beta. Implications for the regulation of cytochrome p450 2C11. J Biol Chem, 1997. 272(30): p. 18718–24.

41. McInnes, L., J. Healy, and J. Melville, Umap: Uniform manifold approximation and projection for dimension reduction. arXiv preprint arXiv:1802.03426, 2018.

42. Stevens, H.Y., et al., Mesenchymal Stromal Cell (MSC) Functional Analysis—Macrophage Activation and Polarization Assays. Bio-protocol, 2024. 14(6).

43. Rui, K., et al., Differential Phase Contrast Imaging to Predict MSC Immune Function. Advanced Healthcare Materials. n/a(n/a): p. 2501553.

44. Ogle, M.E., et al., Hydrogel culture surface stiffness modulates mesenchymal stromal cell secretome and alters senescence. Tissue Engineering Part A, 2020. 26(23-24): p. 1259–1271.

45. Chen, R., et al., Sphingosine 1-phosphate promotes mesenchymal stem cell-mediated cardioprotection against myocardial infarction via ERK1/2-MMP-9 and Akt signaling axis. Life Sciences, 2018. 215: p. 31–42.

46. Zemirli, N., E. Morel, and D. Molino, Mitochondrial dynamics in basal and stressful conditions. International journal of molecular sciences, 2018. 19(2): p. 564.

47. Chiappa, N.A., N. Lal, and E.A. Botchwey, Resolving vs. Non-resolving Sphingolipid Dynamics During Macrophage Activation: A Time-resolved Metabolic Analysis. bioRxiv, 2025: p. 2024.12.31.630925.

48. Yang, K., et al., SGMS1 facilitates osteogenic differentiation of MSCs and strengthens osteogenesis-angiogenesis coupling by modulating Cer/PP2A/Akt pathway. iScience, 2024. 27(4): p. 109358.

49. Fu, P., et al., Nuclear lipid mediators: Role of nuclear sphingolipids and sphingosine-1-phosphate signaling in epigenetic regulation of inflammation and gene expression. J Cell Biochem, 2018. 119(8): p. 6337–6353.

50. Bozzini, N., et al., Epigenetic Regulation Mediated by Sphingolipids in Cancer. Int J Mol Sci, 2023. 24(6).

51. Urs, A.N., E. Dammer, and M.B. Sewer, Sphingosine regulates the transcription of CYP17 by binding to steroidogenic factor-1. Endocrinology, 2006. 147(11): p. 5249–5258.

52. Lucki, N.C. and M.B. Sewer, Nuclear sphingolipid metabolism. Annu Rev Physiol, 2012. 74: p. 131–51.

53. Larey, A.M., et al., High throughput screening of mesenchymal stromal cell morphological response to inflammatory signals for bioreactor-based manufacturing of extracellular vesicles that modulate microglia. Bioactive Materials, 2024. 37: p. 153–171.

